# Therapeutic Efficacy and Safety of Lenvatinib for Unresectable Hepatocellular Carcinoma Beyond Progression with Sorafenib

**DOI:** 10.1101/2020.01.03.893800

**Authors:** Tetsu Tomonari, Yasushi Sato, Hironori Tanaka, Takahiro Tanaka, Yasuteru Fujino, Yasuhiro Mitsui, Akihiro Hirao, Tatsuya Taniguchi, Koichi Okamoto, Masahiro Sogabe, Hiroshi Miyamoto, Naoki Muguruma, Harumi Kagiwada, Masashi Kitazawa, Kazuhiko Fukui, Katsuhisa Horimoto, Tetsuji Takayama

## Abstract

**Background & Aims:** The efficacy and safety of lenvatinib (LEN) as a second/third-line treatment for unresectable hepatocellular carcinoma (HCC) after sorafenib (SOR) therapy remains unknown. We evaluated the outcomes of second/third-line treatment of LEN, investigated the sensitivity of SOR-resistant HCC cell line (PLC/PRF5-R2) to LEN, and their signal transduction pathway by protein array analysis.

**Methods:** We retrospectively enrolled 57 unresectable HCC patients. Radiologic responses in 53 patients were evaluated by modified Response Evaluation Criteria in Solid Tumors. Active signal transduction pathways in cells were identified by protein array analysis, including 1205 proteins.

**Results:** Patients comprised 34 tyrosine kinase inhibitor (TKI)-naive (first-line), nine SOR-intolerant (second-line), and ten resistant to regorafenib (third-line). Objective response rates (ORRs) were 61.8% (21/34) in TKI-naive, 33.3% (3/9) in second-line, and 20.0% (2/10) in third-line groups. The overall survival (OS) and the progression free survival (PFS) in the first-line was significantly longer than those in third-line group (*p*<0.05). Patients with better liver functional reserve (Child score, ALBI grade) exhibited higher ORR and longer OS. LEN was well-tolerated as second/third-line treatment. The IC_50_ value of LEN against PLC/PRF5-R2 cells (30 μM) was significantly higher than that against PLC/PRF5 cells (6.4 μM). LEN inhibited significantly more signal transduction pathways related to FRS2, a crucial FGFR downstream molecule, in PLC/PRF5 than PLC/PRF5-R2 cells.

**Conclusions:** LEN was active and safe as a second/third-line treatment for unresectable HCC. LEN seems to be more effective for HCC patients with better hepatic reserve function or before TKI-resistance is acquired because of partial cross-resistance to SOR.

## Introduction

Hepatocellular carcinoma (HCC) is reportedly the fifth most commonly-diagnosed malignancy and the second leading cause of cancer-related death worldwide.**[1]** For patients with unresectable advanced HCC, sorafenib (SOR) was the first recommended systemic therapy to demonstrate a survival benefit with an adequate safety profile.**[2, 3]** SOR is an oral tyrosine-kinase inhibitor (TKI) that blocks RAF kinase, vascular endothelial growth factor (VEGF) receptors, and the platelet-derived growth factor (PDGF) receptors KIT and FLT3. A phase III SHARP trial showed a median overall survival (mOS) of 10.7 months and a disease control rate (DCR) of 43% in the SOR treatment group of unresectable HCC patients with well-preserved liver function. However, the benefits of SOR were not sustained as the median time-to-progression (mTTP) was only 5.5 months. Subsequently, a randomized, placebo-controlled, phase III RESORCE trial reported that regorafenib (REG), an oral TKI, resulted in survival benefits for patients with advanced HCC who were progressing while on SOR. In this trial, REG showed a 2.8-month improvement in mOS, with a 38% reduction in the risk of death.**[4]** Thus, REG has been the standard second-line chemotherapy for patients refractory to SOR. However, the frequency of advanced HCC patients for whom REG is indicated is reportedly only 30.6‒37%,**[5–8]** and more than half of these patients were not able to receive second-line treatment.

A recent phase III REFLECT trial indicated that lenvatinib (LEN) was non-inferior to SOR as a first-line treatment for unresectable HCC.**[9]** LEN is an oral TKI that targets VEGF receptors 1–3, FGF receptors 1–4, PDGF receptor α, RET, and KIT.**[10–14]** The REFLECT trail showed a mOS of 13.6 months and an mTTP of 8.9 months, where the objective response rate (ORR) was 40.6% in patients of the LEN group. Thus, LEN has been approved in Japan and other countries as a first-line systemic treatment for patients with unresectable advanced HCC.**[9]** Due to the promising efficacy, tolerability, and cost-effectiveness of LEN,**[15]** it has been used not only as a first-line treatment but also as a second-line treatment for patients intolerant to SOR and as a third-line treatment following SOR and REG failure in clinical practice. However, there have only been a few reports regarding the efficacy and adverse effects of LEN when used as a second- or third-line treatment for advanced HCC.**[16, 17]** Especially, little is known about the clinical characteristics of HCC patients that receive potential therapeutic benefits from second- or third-line LEN treatment.

Currently it is not evident whether LEN or SOR should be used as the first-line therapy for advanced HCC. However, both drugs are similar TKIs and some components of the target molecules (VEGFR, PDGFR, KIT) are common to both agents. Therefore, it is highly plausible that they might generate cross resistance to each other. In this context, it is expected that the efficacy of LEN as a second-line or third-line treatment for HCCs, beyond SOR, could be less than that of LEN as a first-line treatment. Moreover, it is unclear which signal transduction pathways are associated with the efficacy of LEN against HCC cells that acquire SOR resistance.

Accordingly, we evaluated the characteristics, therapeutic efficacy, and safety of LEN as a second- and third-line treatment and also as a first-line treatment for unresectable HCC patients in clinical practice. Moreover, to expand upon these clinical findings *in vitro*, we assessed the anti-tumor activity of LEN on a SOR-resistant cell line and performed a comprehensive phosphorylated protein array analysis associated with 377 signal transduction pathways using SOR-resistant and parental HCC cells.

## Material and Methods

### Patient selection and diagnosis of HCC

This retrospective, observational study evaluated the efficacy and safety of LEN (Lenvima®, Eisai Co., Ltd., Tokyo, Japan) monotherapy in patients with unresectable advanced HCC at Tokushima University Hospital between March and December 2018. This study was approved by the Ethics Committee of Tokushima University Hospital (Approval number; 3489). The inclusion criteria were based on those of the REFLECT trial. Briefly, eligible patients had target lesions defined as measurable based on modified Response Evaluation Criteria in Solid Tumors (mRECIST),**[18]** an Eastern Cooperative Oncology Group performance status (ECOG PS) score of 0 or 1,**[19]** Barcelona Clinic Liver Cancer stages (BCLC) B or C categorizations,**[20]** and Child–Pugh class A. Written informed consent was obtained from all patients. The diagnosis of HCC was based on guidelines established by the Liver Cancer Study Group of Japan.**[21]** According to these guidelines, a diagnosis of HCC is confirmed via histology or characteristic radiologic findings such as typical arterial enhancement of the tumor followed by a washout pattern in the images of the portal venous phase or the equilibrium phase obtained by dynamic spiral computed tomography (CT) imaging or contrast-enhanced magnetic resonance imaging (MRI).

### Treatment with LEN

The initial daily oral doses of LEN given to patients weighing ≥ 60, < 60, and < 40 kg were 12, 8, and 4 mg/day, respectively. When serious adverse effects (AEs) were observed, LEN administration was discontinued. Dose interruptions were in accordance with medical package inserts for administering LEN. Briefly, when Grade3 AEs or unacceptable Grade2 AEs developed, LEN was discontinued until AEs recovered and reverted to a lower grade.

### Hepatic reserve function

Hepatic reserve function was assessed according to ALBI grading and Child-Pugh classification. ALBI grade was calculated based on serum albumin and total bilirubin values using the following formula: [ALBI score = (log_10_ bilirubin (µmol/L) × 0.66) + (albumin (g/L) × −0.085)] and defined by the following scores: ≤ −2.60 = Grade 1, > −2.60 to ≤ −1.39 = Grade 2, > −1.39 = Grade 3.**[22]**

### Follow-up and patient outcome

Patients were observed for at least 12 weeks. Safety was assessed by recording any adverse drug reactions, clinical laboratory tests, physical examination, measurement of vital signs, hematological and biochemical laboratory testing, and urinalysis. Adverse drug reactions were defined according to the Common Terminology Criteria for Adverse Events version 5.0. Radiologic responses to therapy were evaluated according to mRECIST at the 8th week after starting LEN and every 8 weeks thereafter. ORR was defined as the sum of complete response (CR) and partial response (PR) rates. DCR was defined as the sum of CR, PR, and stable disease (SD) rates. Progression free survival (PFS) was defined as the time from the first day of administering LEN until the day of radiological progression.

### Cell culture and viability analysis

The representative human hepatoma cell line PLC/PRF5 was purchased from the American Tissue Culture Collection (ATCC, Manassas, VA). Establishment of the SOR-resistant PLC/PRF5 cell line (PLC/PRF5-R2) was performed as described previously.**[23]** Cells were grown in Dulbecco’s modified Eagle’s medium (Invitrogen Sigma-Aldrich, St. Louis, MO) supplemented with 10% FBS and 2 mM L-glutamine. Cell viability was assessed via a 3-(4, 5-dimethylthiazol-2-yl)-2,5-diphenyl-2H-tetrazolium bromide (MTT) assay as described previously.**[23]**

### Identification of active signal transduction pathways by protein array

We used a self-made comprehensive protein phosphorylation array that included 1205 proteins representing 377 pathways involved in signal transduction, as described in the supporting information (Fig S1) to determine the phosphorylation status of selected proteins in the active signal transduction pathways of HCC cells in the absence or presence of LEN. The difference in the degree of phosphorylation of each pathway between the two groups was estimated by Welch’s t-test (statistical significance: *p* < 0.05), and the number of proteins showing significant probability was counted for each pathway. The probability of each pathway was then estimated based on the hyper-geometric distribution of the 377 pathways comprising the 1205 proteins.

### Statistical analysis

Categorical variables were compared using the Fischer’s exact test, whereas continuous variables were compared using Mann-Whitney and Kruskal-Walls tests. All significance tests were two-tailed, and statistical significance was set at *p* < 0.05. Kaplan–Meier plots of medians (with 95% confidence interval, [CI])) were used to estimate PFS. All statistical analyses were undertaken using Easy R (EZR) version 1.29 (Saitama Medical Center, Jichi Medical University, Saitama, Japan).

## Results

### Patient characteristics

A total of 57 patients with unresectable HCC who had received LEN were enrolled in this study. However, of these, 4 patients were excluded from the analysis because they could not be evaluated using mRECIST measurements due to renal failure. Thus, 53 patients were retrospectively analyzed. Baseline characteristics of these patients are listed in Table 1. The median observation period following the initiation of treatment with LEN was 266 (111–603) days. The median age of the patients was 71 years (range, 47–85 years). Of all patients, 14 (26.4%) were HBV antigen-positive and 22 (41.5%) were HCV antibody-positive. The ECOG PS was 0 in 48 patients (90.6%). The median AFP value was 37 ng/ml (range 2–568100) and Child-Pugh scores before treatment were 5 points in 30 patients and 6 points in 23 patients. ALBI grades before treatment were 1 point in 22 patients and 2 points in 31 patients. LEN therapy was initiated at BCLC stage B in 37 patients and at stage C in 16 patients. The median number of cases of transarterial chemoembolization (TACE) before treatment with LEN was 1 (0‒9). Among the 53 patients, 34 were TKI-naive (first-line), nine were intolerant to SOR (second-line), and ten were resistant to SOR and REG (third-line). The median duration of follow-up in each group was as follows: 255 (118–603) days for first-line, 391 (111–603) days for second-line, and 265 (132–507) days for third-line. There were no significant differences in patient characteristics between those with and without a previous history of TKI treatment, including hepatic reserve function and tumor burden.

**Table 1.** Characteristics of patients with unresectable hepatocellular carcinoma treated with lenvatinib.

| Characteristics | All<br>(*n* = 53) | First line<br>(*n* = 34) | Second line<br>(*n* = 9) | Third line<br>(*n* = 10) | *p*-<br>value |
| --- | --- | --- | --- | --- | --- |
| Age, median<br>[range], (years) | 71<br>[47-85] | 72<br>[53-85] | 70<br>[61-80] | 66<br>[47-83] | 0.45 |
| Gender (male/female), *n* | 44/9 | 30/4 | 7/2 | 7/3 | 0.38 |
| Etiology (HBV/HCV/NBNC), *n* | 14/22/17 | 10/12/12 | 2/4/3 | 2/6/2 | 0.87 |
| ECOG PS (0/1), *n* | 48/5 | 32/2 | 7/2 | 10/0 | 0.24 |
| Platelets, median<br>[range], (10<sup>4</sup>/μl) | 14.4<br>[6.2-31.8] | 14.4<br>[6.2-31.8] | 16.3<br>[9.0-25.7] | 11.4<br>[6.2-23.7] | 0.45 |
| M2BpGi<br>[range] (C.O.I) | 1.54<br>[0.44-13.1] | 1.25<br>[0.48-5.76] | 1.59<br>[0.44-13.1] | 2.23<br>[0.58-5.4] | 0.17 |
| Child-Pugh score (5/6/7/8), *n* | 30/23/0/0 | 22/12/0/0 | 5/4/0/0 | 3/7/0/0 | 0.15 |
| ALBI Grade (1/2/3), *n* | 22/31/0 | 13/21/0 | 4/5/0 | 1/9/0 | 0.08 |
| Number of intrahepatic lesions<br>(None/1/2-7/>7) | 0/6/23/24 | 0/4/17/13 | 0/1/4/4 | 0/1/2/7 | 0.70 |
| Maximum size of intrahepatic<br>lesion (None/≤50/>50) (mm) | 0/42/11 | 0/26/8 | 0/7/2 | 0/9/1 | 0.79 |
| Portal vein invasion<br>(absent/present), *n* | 47/6 | 30/4 | 7/2 | 9/1 | 0.72 |
| Extrahepatic spread<br>(absent/present), *n* | 42/11 | 29/5 | 5/4 | 8/2 | 0.16 |
| AFP, median<br>[range] (ng/ml) | 37<br>[2-568100] | 12<br>[2-568100] | 414<br>[4-2262] | 37<br>[4-5050] | 0.73 |
| BCLC stage (B/C), *n* | 37/16 | 24/10 | 5/4 | 8/2 | 0.51 |
| Previous treatment times of<br>TAE/TACE [range] | 1<br>[0-9] | 1<br>[1-6] | 2<br>[0-9] | 2<br>[1-6] | 0.10 |
| Initial dose of Lenvatinib<br>(12/8/4), (mg), *n* | 30/22/1 | 23/10/1 | 3/6/0 | 4/6/0 | 0.13 |

HBV: hepatitis B virus, HCV: hepatitis C virus, NBNC: non B non C, ECOG PS: Eastern Cooperative Oncology Group performance status, M2BPGi mac-2 binding protein glycosylation isomer, ALBI: albumin-bilirubin, AFP: alpha- fetoprotein, BCLC: Barcelona Clinic Liver Cancer, TAE/TACE: transcatheter embolization/chemoembolization.

### Response to LEN

Fifty-three patients had measurable lesions that could be evaluated by enhanced CT/MRI at 8 weeks after the initiation of LEN treatment. Of these 53 patients, two exhibited a CR (3.8%), 24 had a PR (45.3%), 25 had SD (47.2%), and two presented with progressive disease (PD) (3.8%). The ORR and DCR were 49.1% (26/53) and 96.2% (51/53), respectively (Table S1). Regarding the response in each treatment-line group, ORRs in the first-line group (61.8%; 21/34) were higher than those in the second-line group (33.3%, 3/9; *p* = 0.28) and were higher than those in the third-line group (20.0%, 2/10; *p* = 0.27; Table 2). Moreover, the ORR with BCLC stage B (20/37, 54.1%) was higher than that with BCLC stage C (6/16, 37.5%). In terms of hepatic reserve functions, the ORR in the Child -Pugh score of 5 group (16/30, 53.3%) was higher than that with a Child-Pugh score of 6 (10/23, 43.5%). Likewise, ORR in the ALBI grade 1 group (14/22, 58.8%) was higher than that in the ALBI grade 2 group (12/31; 38.7%; Table 2). The median PFS of the 53 patients was 8.5 months (95% CI: 6.9–13.8 months; Fig S2). The PFS in the first-line group was significantly longer than that in the second-line group (*p* < 0.05; Fig 1a). The PFS in the first-line group was significantly longer than that in the third-line group (*p* < 0.01). The PFS in the BCLC stage B group was tended to be longer than that in the stage C group (*p* = 0.07; Fig 1b). Similarly, PFS in the ALBI Grade1 group was significantly longer than that in the ALBI Grade2 group (*p* < 0.05; Fig 1c). Further, PFS in cases with a Child-Pugh score of 5 was significantly longer than that in cases with a Child-Pugh score of 6 (*p* < 0.01; Fig 1d).

**Table 2.** Response to treatment with lenvatinib for advanced hepatocellular carcinoma according to. treatment line, stage, and hepatic functional reserve

| Evaluation (mRECIST) | CR | PR | SD | PD | ORR (%) | DCR (%) |
| --- | --- | --- | --- | --- | --- | --- |
| Treatment line |  |  |  |  |  |  |
| First line ($n = 34$) | 1 (2.9) | 20 (58.8) | 12 (35.3) | 1 (2.9) | 61.8 | 97.1 |
| Second line ($n = 9$) | 1 (11.1) | 2 (22.2) | 5 (55.5) | 1 (14.3) | 33.3 | 88.8 |
| Third line ($n = 10$) | 0 (0) | 2 (20.0) | 8 (80.0) | 0 (0) | 20.0 | 100 |
| BCLC stage |  |  |  |  |  |  |
| B ($n = 37$) | 2 (5.4) | 18 (48.6) | 17 (45.9) | 0 (0) | 54.1 | 100 |
| C ($n = 16$) | 0 (0) | 6 (37.5) | 8 (50.0) | 2 (12.5) | 37.5 | 87.5 |
| Child–Pugh score |  |  |  |  |  |  |
| 5 ($n = 30$) | 2 (6.7) | 14 (46.7) | 12 (40.0) | 2 (6.7) | 53.3 | 93.3 |
| 6 ($n = 23$) | 0 (0) | 10 (43.5) | 13 (56.5) | 0 (0) | 43.5 | 100 |
| ALBI grade |  |  |  |  |  |  |
| 1 ($n = 22$) | 1 (4.5) | 13 (59.0) | 6 (27.2) | 2 (9.1) | 63.6 | 91.0 |
| 2 ($n = 31$) | 1 (3.2) | 11 (35.5) | 19 (61.3) | 0 (0) | 38.7 | 100 |

*mRECIST* modified response evaluation criteria in solid tumors, *BCLC* barcelona clinic liver cancer, *ALBI* albumin-bilirubin, *CR* complete response, *PR* partial response, *SD* stable disease, *PD* progressive disease, *ORR* overall response rate, *DCR* disease control rate.

**Figure 1.**
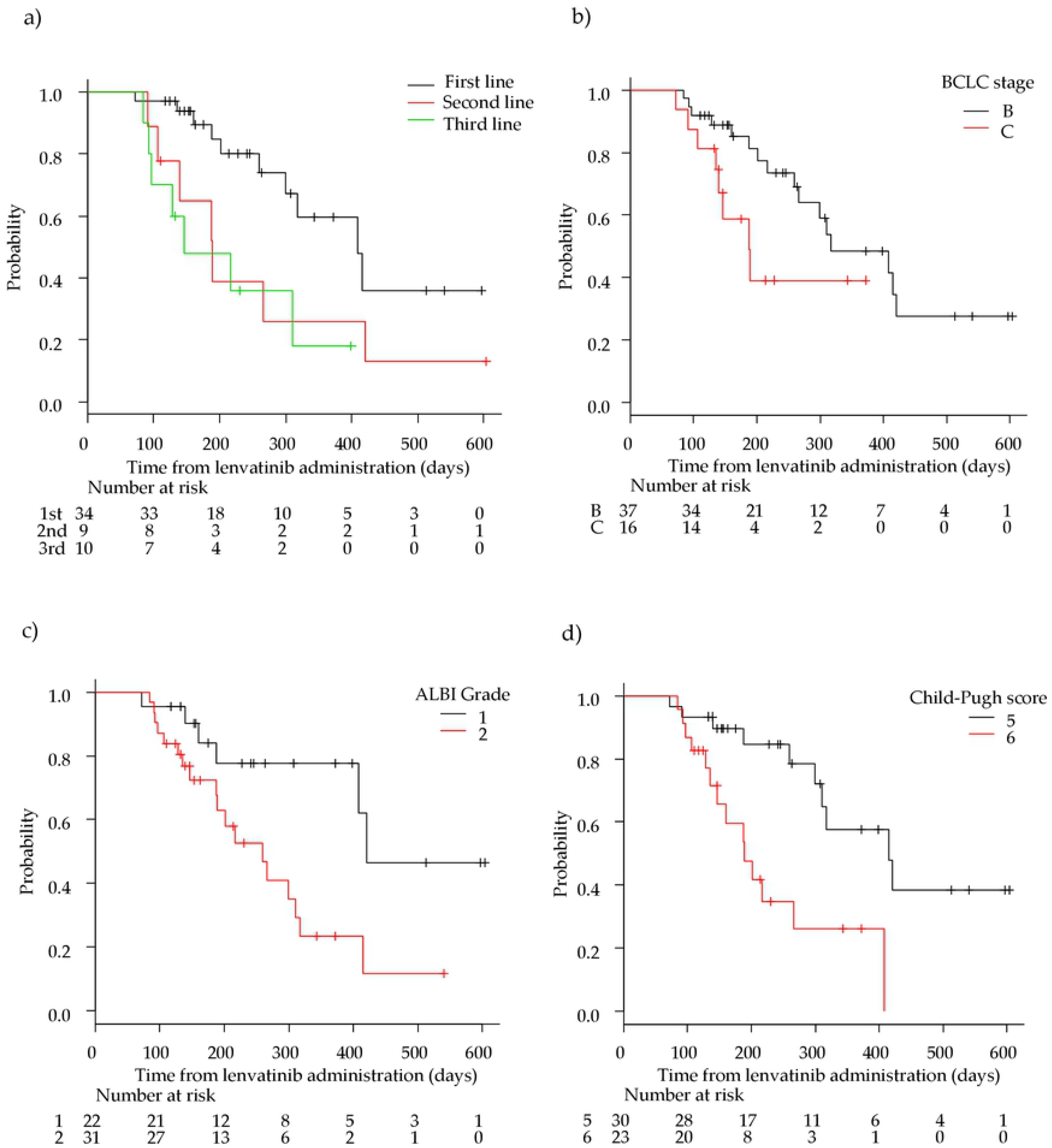
Kaplan–Meier analysis of progression free survival among patients with advanced hepatocellular carcinoma treated with lenvatinib according to treatment lines and hepatic functional reserve. **a** First/second/third-line groups. **b** Barcelona clinic liver cancer (BCLC) stage B and C groups. **c** Albumin-bilirubin (ALBI) grade 1 and 2 group **d** Child-Pugh score 5 and 6 groups.

The median OS of the 53 patients was NA (95% CI:19.8–NA month; Fig S3). The median OS in the first-line group, in the second-line and in the third group were not reached. The OS in the first-line group was significantly longer than that in the third-line group (*p* < 0.05; Fig 2a). There was no significant difference in OS between the first-line and second-line groups. The OS in the BCLC stage B group was significantly longer than that in the stage C group (*p* < 0.01; Fig 2b). The OS in the ALBI Grade1 group tended to be longer than that in the ALBI Grade2 group (*p* < 0.05; Fig 2c). Moreover, the OS with a Child-Pugh score of 5 was significantly longer than that with a score of 6 (*p* < 0.05; Fig 2d).

**Figure 2.**
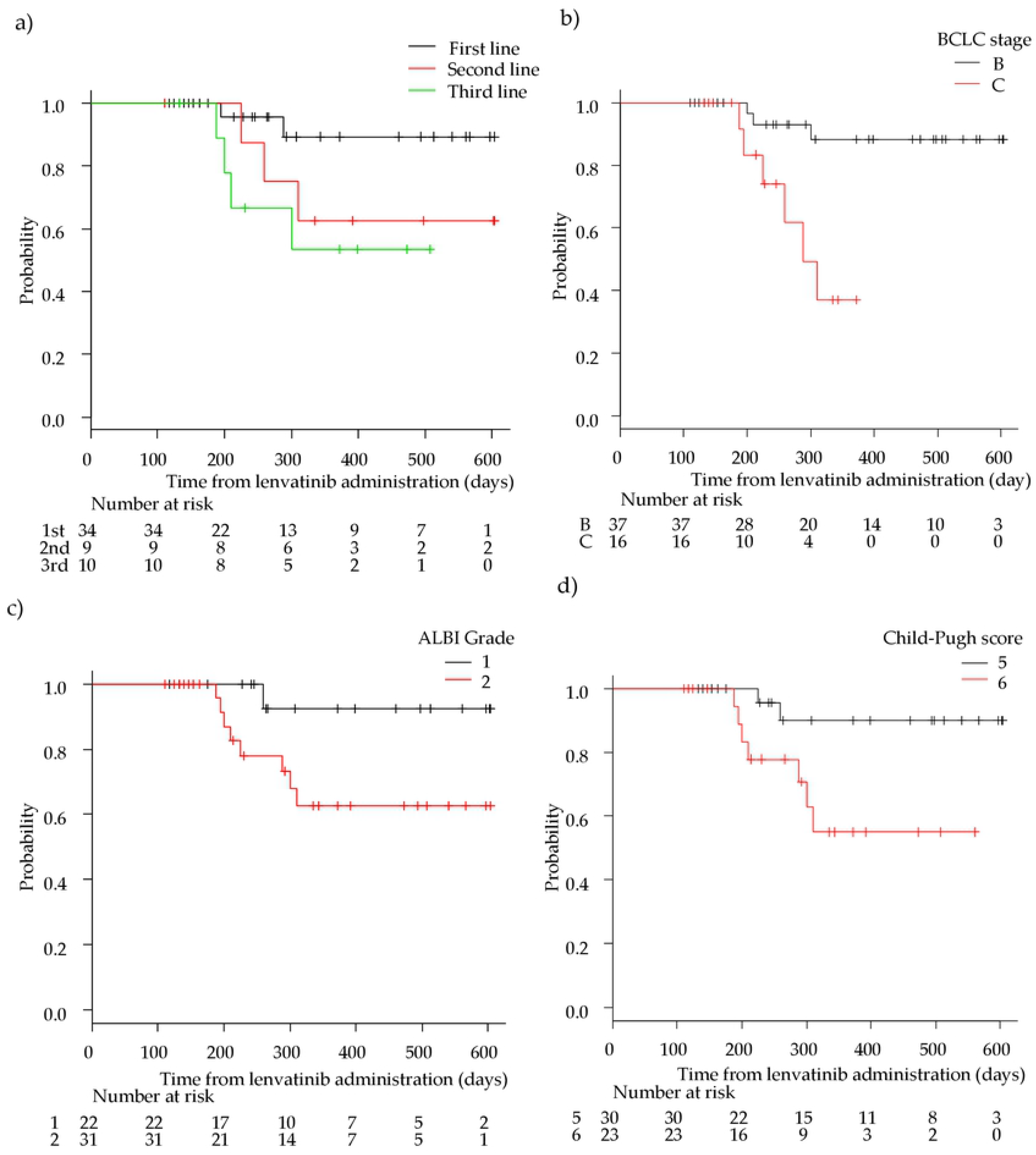
Kaplan–Meier analysis of overall survival among patients with advanced hepatocellular carcinoma treated with lenvatinib according to treatment lines and hepatic functional reserve. **a** First/second/third-line groups. **b** Barcelona clinic liver cancer (BCLC) stage B and C groups. **c** Albumin-bilirubin (ALBI) grade 1 and 2 group **d** Child-Pugh score 5 and 6 groups.

### Safety

Grade 4 AEs were not observed during the observation period. The most common all-grade drug-related AEs were hypertension (54.7%; 29/53), proteinuria (47.2%; 25/53), fatigue (49.1%; 26/53), appetite loss (37.7%; 20/53), and palmar-plantar erythrodysesthesia (26.4%; 14/53; Table 3). The most common grade 3 drug-related AEs were proteinuria (24.5%, 13/53), hypertension (15.1%, 8/53), fatigue (7.5%, 4/53), diarrhea (3.8%, 2/53). There were no significant differences in LEN-related AEs among each treatment group. Moreover, the frequencies of LEN-related AEs were higher in the ALBI Grade2 group than in the ALBI Grade1 group (Table 4). Among them, the frequency of fatigue was significantly higher in patients in the ALBI-2 group (23/31, 74.2%) than in those in the ALBI-1 group (3/22 13.6%; *p* < 0.01). Similar AE results were observed between groups comprising Child-Pugh scores of 5 and 6 (data not shown). Treatment with LEN was discontinued due to AEs in only three patients. All AEs were controlled by appropriate dose reduction or care.

**Table 3.**
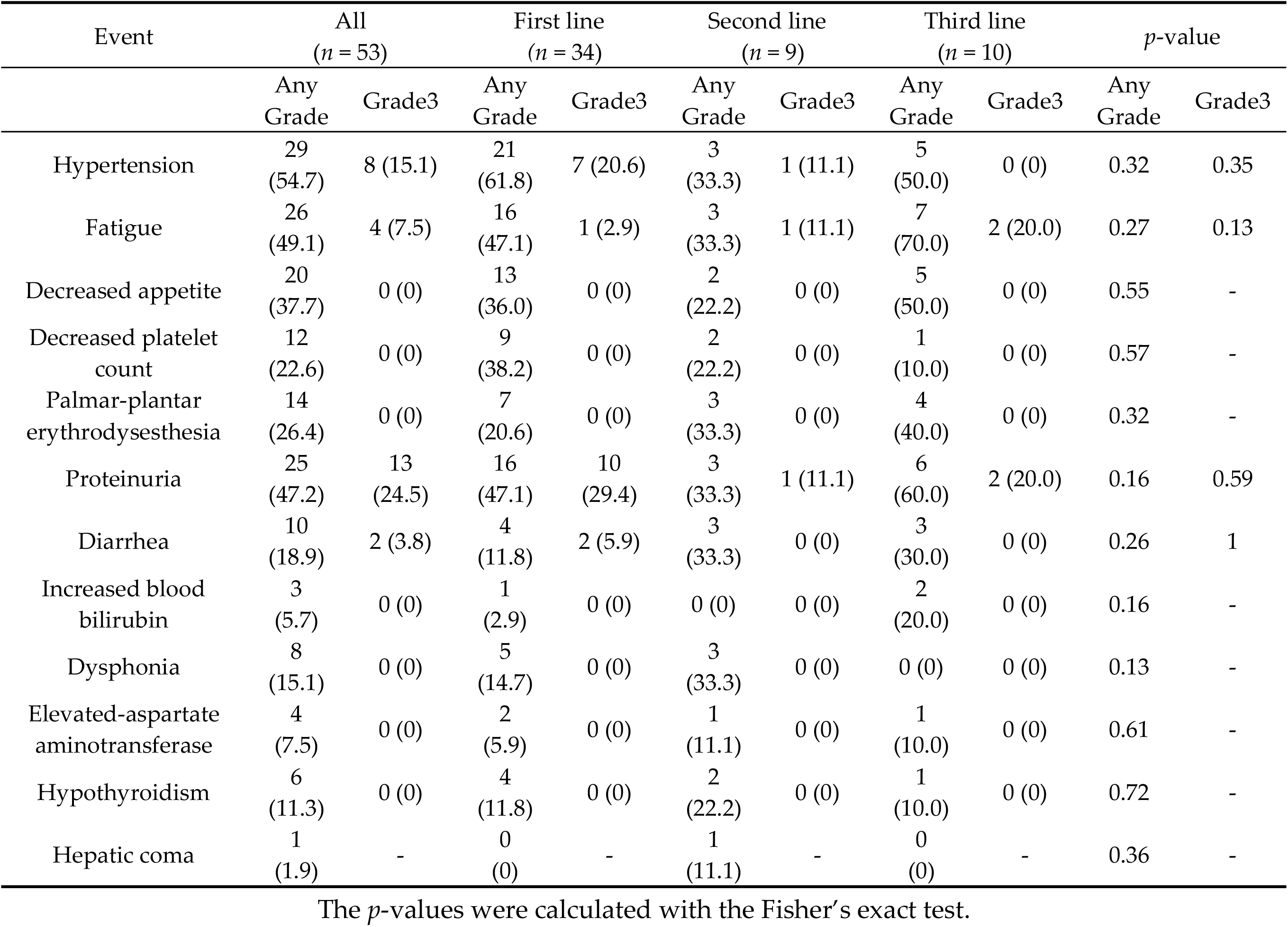
Adverse events of lenvatinib treatment.

| Event | All<br>(*n* = 53) |  | First line<br>(*n* = 34) |  | Second line<br>(*n* = 9) |  | Third line<br>(*n* = 10) |  | *p*-value |  |
| --- | --- | --- | --- | --- | --- | --- | --- | --- | --- | --- |
|  | Any<br>Grade | Grade3 | Any<br>Grade | Grade3 | Any<br>Grade | Grade3 | Any<br>Grade | Grade3 | Any<br>Grade | Grade3 |
| Hypertension | 29<br>(54.7) | 8 (15.1) | 21<br>(61.8) | 7 (20.6) | 3<br>(33.3) | 1 (11.1) | 5<br>(50.0) | 0 (0) | 0.32 | 0.35 |
| Fatigue | 26<br>(49.1) | 4 (7.5) | 16<br>(47.1) | 1 (2.9) | 3<br>(33.3) | 1 (11.1) | 7<br>(70.0) | 2 (20.0) | 0.27 | 0.13 |
| Decreased appetite | 20<br>(37.7) | 0 (0) | 13<br>(36.0) | 0 (0) | 2<br>(22.2) | 0 (0) | 5<br>(50.0) | 0 (0) | 0.55 | - |
| Decreased platelet<br>count | 12<br>(22.6) | 0 (0) | 9<br>(38.2) | 0 (0) | 2<br>(22.2) | 0 (0) | 1<br>(10.0) | 0 (0) | 0.57 | - |
| Palmar-plantar<br>erythrodysesthesia | 14<br>(26.4) | 0 (0) | 7<br>(20.6) | 0 (0) | 3<br>(33.3) | 0 (0) | 4<br>(40.0) | 0 (0) | 0.32 | - |
| Proteinuria | 25<br>(47.2) | 13<br>(24.5) | 16<br>(47.1) | 10<br>(29.4) | 3<br>(33.3) | 1 (11.1) | 6<br>(60.0) | 2 (20.0) | 0.16 | 0.59 |
| Diarrhea | 10<br>(18.9) | 2 (3.8) | 4<br>(11.8) | 2 (5.9) | 3<br>(33.3) | 0 (0) | 3<br>(30.0) | 0 (0) | 0.26 | 1 |
| Increased blood<br>bilirubin | 3<br>(5.7) | 0 (0) | 1<br>(2.9) | 0 (0) | 0 (0) | 0 (0) | 2<br>(20.0) | 0 (0) | 0.16 | - |
| Dysphonia | 8<br>(15.1) | 0 (0) | 5<br>(14.7) | 0 (0) | 3<br>(33.3) | 0 (0) | 0 (0) | 0 (0) | 0.13 | - |
| Elevated-aspartate<br>aminotransferase | 4<br>(7.5) | 0 (0) | 2<br>(5.9) | 0 (0) | 1<br>(11.1) | 0 (0) | 1<br>(10.0) | 0 (0) | 0.61 | - |
| Hypothyroidism | 6<br>(11.3) | 0 (0) | 4<br>(11.8) | 0 (0) | 2<br>(22.2) | 0 (0) | 1<br>(10.0) | 0 (0) | 0.72 | - |
| Hepatic coma | 1<br>(1.9) | - | 0<br>(0) | - | 1<br>(11.1) | - | 0<br>(0) | - | 0.36 | - |

The *p*-values were calculated with the Fisher's exact test.

**Table 4.** The relationship between adverse events and ALBI-grade.

| Event | ALBI grade *n* = 53 |  | *p*-value |
| --- | --- | --- | --- |
|  | ALBI-1<br>*n* = 22 | ALBI-2<br>*n* = 31 |  |
| Hypertension | 11 (50.0) | 18 (58.1) | 0.59 |
| Fatigue | 3 (13.6) | 23 (74.2) | < 0.01 |
| Decreased appetite | 7 (31.8) | 13 (41.9) | 0.57 |
| Decreased platelet count | 4 (18.2) | 8 (25.8) | 0.74 |
| Palmar-plantar erythrodysesthesia | 5 (22.7) | 9 (29.0) | 0.76 |
| Proteinuria | 10 (45.5) | 15 (48.4) | 1 |

ALBI albumin-bilirubin; the *p*-values were calculated with the Fisher's exact test.

### Drug administration

The relative dose intensity (RDI) in the first-, second-, and third-line groups was 83.1%, 73.6%, and 73.5%, respectively. Treatment continued for 12 weeks in all cases, except in one case of PD and three cases of withdrawal due to AEs (two cases of grade 3 fatigue). During 12 weeks of observation, AEs led to interruption of the administration of LEN in eight (15.0%) patients and dose reduction in 13 (24.5%).

### In vitro cell viability of the SOR-resistant cell line after LEN treatment

To confirm our clinical observations and to analyze mechanisms underlying the sensitivity of HCC cells to LEN *in vitro*, we first performed a cell viability assay using previously established PLC/PRF5 and SOR-resistant PLC/PRF5-R2 cell lines.[23] The IC_50_ value of LEN towards PLC/PRF5 cells was 6.4 µM, and this value was consistent with those of previous reports.[23] However, for SOR-resistant PLC/PRF5-R2 cells, the IC_50_ value of LEN was 30 µM, which was significantly higher than that with the parental PLC/PRF5 cells (*p* < 0.05; Fig 3A). These findings suggested that PLC/PRF5-R2 cells might show partial cross-resistance to LEN.

**Figure 3.**
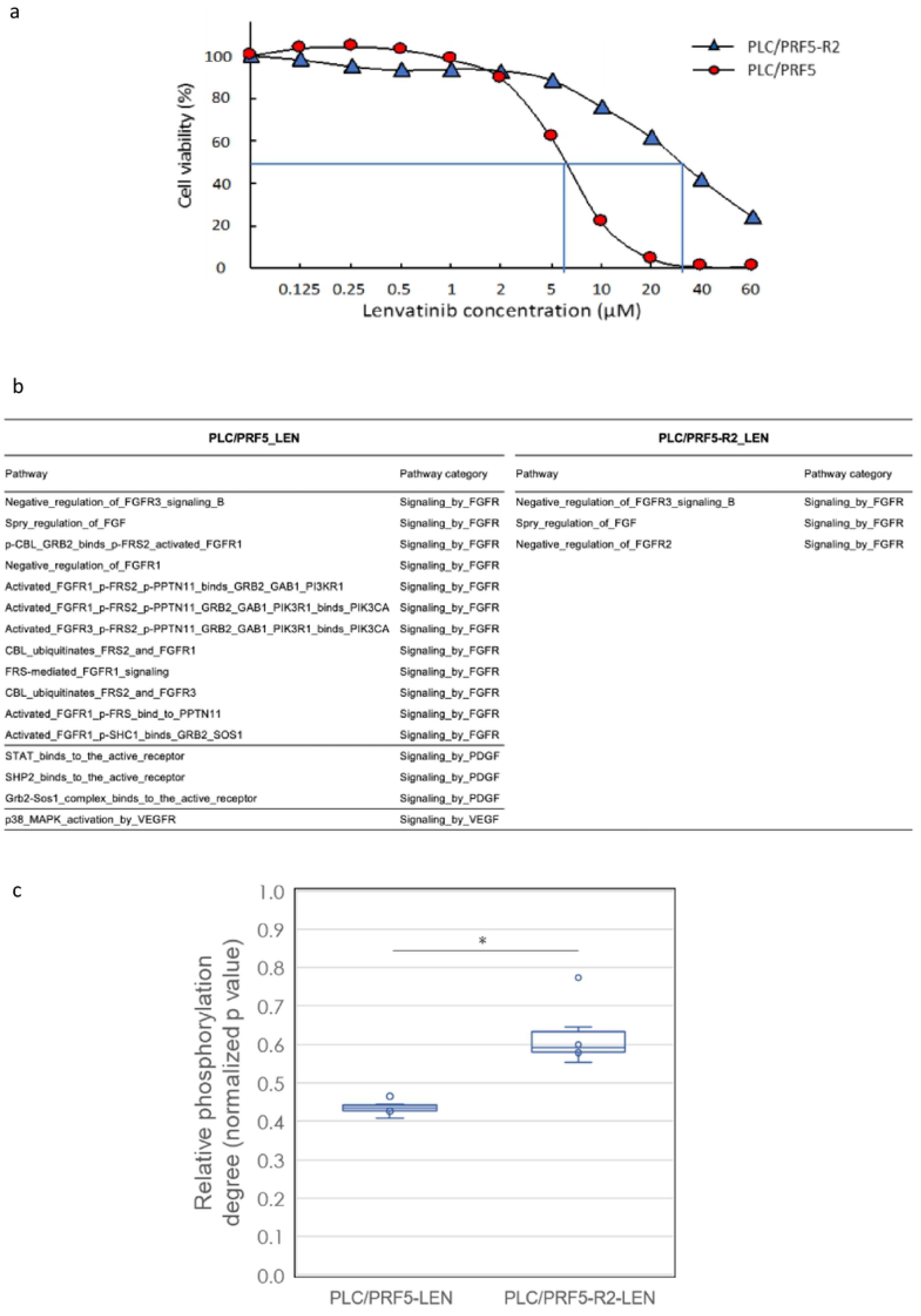
*In vitro* cell viability and signal transduction pathway analysis of SOR resistant cell line with lenvatinib treatment by a comprehensive protein phosphorylation array. **a** The sensitivities of PLC/PRF5 and PLC/ PRF5-R2 to lenvatinib (LEN) were assessed by MTT assay. **b** A list of LEN-related signal transduction pathways significantly altered after treatment with LEN in PLC/PRF5 and PLC/PRF5-R2 cells. The 114 LEN related pathways were categorized into “Signaling by FGFR”, “Signaling by PDGF”, “Signaling by VEGFR”, and “Signaling by SRC-KIT” among 377 pathways tested. **c** Boxplot analysis of degree of phosphorylation of associated proteins belonging to the 63 FRS2-related transduction pathways in PLC/PRF5 and PLC/ PRF5-R2 cells in response to LEN. The boxes show the interquartile rage with the median value indicated by the horizontal line, whiskers show the range, circles indicate outliers. *p<0.0014.

### In vitro signal transduction pathway analysis in SOR-resistant cell lines

LEN reportedly inhibits the phosphorylation of tyrosine-kinases such as FGF receptors, VEGF receptors, and the PDGF receptors RET and KIT.[10–13] Therefore, we investigated the degree of protein phosphorylation related to LEN-related signal transduction pathways in response to LEN using a comprehensive protein phosphorylation array (Fig S1). A representative heatmap of array results including cluster analysis is shown in Fig S4. The heatmap demonstrated that protein phosphorylation levels in all 377 signaling pathways were clearly distinguished and clustered between PLC/PRF5 and PLC/PRF5-R2 cells following treatment with LEN. The number of LEN-related signal transduction pathways that were significantly altered following LEN treatment was 16 including 12 related to FGFR, three to PDGF, and one to VEGF, whereas only three FGFR pathways were altered in PLC/PRF5-R2 cells (Fig 3B). These results further indicated that PLC/PRF5-R2 cells show cross-resistance to LEN.

### In vitro FRS2-related pathway analysis in the SOR-resistant cell line

The adaptor protein fibroblast growth factor receptor substrate 2 (FRS2) is reportedly an essential downstream component of the FGFR signaling pathway, and acts as a hub linking several signaling pathways to ultimately activate FGFRs.[24] Therefore, we examined the inhibitory effects of LEN on the phosphorylation of 63 FRS2-related pathways among a total of 377 pathways involved in signal transduction (Table S2) in PLC/PRF5 and PLC/PRF5-R2 cell lines using a phosphorylation array (Fig 3C). LEN suppressed the phosphorylation of FRS2 protein in those pathways in both cell lines. However, the degree of suppression was significantly higher in PLC/PRF5 cells than in PLC/PRF5-R2 cells (*p* < 0.01, Welch’s t-test); this indicated that the degree of LEN-mediated inhibition of FRS2-related signaling pathways in PLC/PRF5-R2 cells was significantly lower than that in PLC/PRF5 cells. Thus, it was evident that SOR-resistant HCC cells show partial cross-resistance to LEN based on resistance to the LEN-mediated inhibition of FGFR signaling pathways.

## Discussion

In this retrospective study, we demonstrated the therapeutic efficacy and safety of LEN as a second- and third-line treatment, particularly for patients intolerant to SOR (second-line) and as a first-line treatment for HCC. Moreover, our results suggested that treatment with LEN while maintaining better hepatic functional reserves might exert more beneficial effects on the prognosis of patients with advanced HCC in a clinical setting. Furthermore, our *in vitro* experiments revealed that the SOR-resistant cell line PLC/PRF5-R2 was partially cross-resistant to LEN. LEN significantly inhibited 16 signal transduction pathways including 12 FGFR pathways in parent PLC/PRF5 cells but inhibited only a few pathways in PLC/PRF5-R2 cells. These results support our clinical observations indicating that the response rate of third-line LEN treatment was rather low compared to that of first-line treatments.

REG has been used only to treat patients who are tolerant to SOR treatment, as a second-line treatment following SOR, but not for patients intolerant to SOR. Approximately one third of patients who receive SOR treatment are reportedly intolerant to SOR,[5–8] and therefore are deemed unsuitable to receive substantially effective systemic chemotherapy. Our data showed a somewhat high ORR (26.3%), as well as a favorable safety profile, for patients intolerant to SOR. Thus, LEN shows potential as a second-line treatment for patients with unresectable HCC intolerant to SOR.

In this study, the ORRs of LEN in third-line treatment (20.0%) were significantly lower than those in first-line treatment group (61.8%) and somewhat lower than those in second-line treatment group (33.3%). However, DCR in the third-line treatment group was highly similar to that with the other-line treatments. Likewise, the median PFS and OS in the third-line treatment group were also significantly shorter than those in the first-line treatment group and tended to be shorter than those in the second-line treatment groups.

To date, there have been no studies investigating ORRs for second- and third-line treatment regimens using LEN. Hiraoka et al. reported that there were no significant differences in the efficacy of LEN for advanced HCC between TKI-naïve and TKI-experienced groups.[16] Although ORRs for each patient group were not shown, the TKI-experienced patients included only 25% (11/44) of patients following SOR–REG treatment (third line), whereas 75% (33/44) were considered SOR-intolerant patients (second line). In this context, it appears that LEN is relatively effective for SOR-intolerant patients but less effective for patients resistant to SOR–REG treatment. In fact, the effects of LEN in our second-line cohort were comparable to those with first-line treatment. This might be due to the fact that patients administered second-line LEN treatment could still be sensitive to TKIs since they could not continue SOR treatment due to of detrimental adverse effects but did not acquire complete resistance to SOR. Therefore, it is reasonable to assume that LEN is fairly effective for HCC patients intolerant to SOR as a second-line treatment.

Our *in vitro* cell viability assay revealed that the IC_50_ of LEN towards the SOR-resistant cell line (PLC/PRF5-R2) was significantly higher than that with parental PLC/PRF5 cells (*p* < 0.01; Fig 3A). It is plausible that the signaling pathway inhibited by LEN in HCC cells was modified during the course of acquiring SOR-resistance, leading to partial cross-resistance to LEN. In general, it is difficult to delineate the multifaceted and dynamic pathway regulation in response to TKIs such as LEN. Therefore, we employed a comprehensive protein phosphorylation array, which can simultaneously measure the phosphorylation degrees of 1205 proteins belonging to 377 signal transduction pathways (Fig S1). As a result, we confirmed that LEN mainly inhibited the phosphorylation of 12 FGFR-related pathways in parental PLC/PRF5 cells, consistent with previous reports showing that LEN selectively suppresses the proliferation of HCC cells with activated FGF signaling pathways; this is a distinct feature of LEN as compared to that with SOR.[24] Conversely, only three FGFR-related signaling pathways were significantly inhibited in PLC/PRF5-R2 cells, indicating the partial resistance of those SOR-resistant cells to LEN. These data were further supported by the fact that the phosphorylation degree of FRS2, which plays a pivotal role in FGFR-related pathways, in PLC/PRF5-R2 cells was significantly higher than that in parental PLC/PRF5 cells (Figure 3C). Thus, our protein array analysis suggests that LEN is less effective for HCC patients with resistance to SOR than for SOR-naïve patients due to cross-resistance between LEN and SOR.

The AE profiles in this study were similar to those in previous reports,[16, 17, 25] mostly documented during first-line treatment. Otherwise, the incidence of AEs with second/third line treatment was similar to that with first-line treatment (Table 3). This might be explained by the particular characteristics of LEN, which shows different AE spectra from those of both SOR and REG. Moreover, treatment with LEN could easily be initiated as a second/third-line treatment following treatment with SOR or REG, even in patients suffering from severe AEs related to SOR or REG such as hand–foot syndrome and diarrhea. Thus, our data demonstrated the safety and feasibility of LEN as a second/ third-line treatment for unresectable HCC. In addition, the incidence of fatigue in the ALBI-2 group containing all treatment lines was significantly higher than that in the ALBI-1 group (Table 4). This fatigue was often the cause of dose reduction and the interruption of treatment, especially in the ALBI-2 group. Since the ALBI score is calculated from only albumin and total bilirubin values, the ALBI-2 group is more likely to have lower serum albumin levels, reflecting poorer nutrition and performance status, which might more readily lead to fatigue in HCC patients.[26–28]

In this study, patients with better liver functional reserve (Child-Pugh score of 5, ALBI Grade1) showed better response to LEN and longer OS. Recently, Ueshima and associates analyzed 82 patients with unresectable HCCs treated with LEN and reported that ALBI Grade1 and serum AFP levels < 200 are predictors of high response rate.[29] They also demonstrated that the time to treatment failure in patients with better liver functional reserve was longer. Their patients included 61.0% TKI-naïve, 24.4% second-line (SOR intolerance), and only 14.6% third-line treatment patients. They also included those with a Child-Pugh score B as well as those with score A, and BCLC stages A, B, and C. Although the specific proportions of patients were largely different from those in our study, our data on treatment outcome displayed partial similarity to theirs. Patients with better liver functional reserve showed better response, and the OS in the Child-Pugh score 5 group was significantly longer than that in the Child-Pugh score 6 group. These results might be partly explained by the difference in RDI. The RDI of LEN in the Child-Pugh score 5 group (81.4%) was higher than that in the Child-Pugh score 6 group (76.5%). Thus, to maximize the therapeutic effect of LEN, it should be used in patients with unresectable HCC while liver function is preserved, as with Child-Pugh A and ALBI-1 grade patients. Although repeated TACE was often performed for the treatment of unresectable HCC until recently, LEN treatment should be initiated in HCC patients with better liver functional reserve. Eventually, in this study, patients with BCLC stage C showed lower ORRs and shorter PFS and OS than those with BCLC stage B. One of the reasons is that BCLC stage C cases included six patients with portal vein invasion, which might be related to the poor outcomes observed.

One limitation of this study was its single-center, retrospective design. Another limitation was that the observation period was relatively short and the number of analyzed patients was small. However, considering that LEN had only been approved in Japan for 17 months, our observations at the specified cutoff date are adequate to report real-world treatment results, especially those related to evaluating the initial safety and efficacy of the clinical use of LEN. A large-scale prospective study is indispensable to establish the efficacy of LEN for second-and third line-treatment use in the future. Regarding *in vitro* experiments, LEN exerts its effect by blocking of not only FGFR, but also PDGFR-α and VEGFR, the latter of which is expressed in endothelial cells rather than cancer cells. Therefore, the findings should be confirmed using *in vivo* HCC xenograft models, where the anti-angiogenic activity of LEN can be evaluated.

## Conclusions

In conclusion, we demonstrated that LEN monotherapy could be feasible as a second/third-line treatment for unresectable HCC and suggest that LEN should preferably be applied to patients with better functional liver reserve (ALBI-1 or Child score 5) to obtain good outcomes. Moreover, LEN was more effective in TKI-naïve patients as a first-line treatment than in patients administered LEN for second- and third-line treatment, and particularly in patients on third-line treatment after SOR– REG treatment. These clinical data are supported by *the in vitro* experimental results indicating that the SOR-resistant cell line became partially cross-resistance to LEN by altering FGFR-related signal transduction pathways.

## Author’s contributions

All authors contributed to the study conception and design. Material preparation, data collection and analysis were performed by Tetsu Tomonari, Hironori Tanaka, Takahiro Tanaka, Yasuteru Fujino, Yasuhiro Mitsui, Koichi Okamoto, Hiroshi Miyamoto, Naoki Muguruma, Harumi Kagiwada, Masashi Kitazawa, Kazuhiko Fukui, and Katsuhisa Horimoto. The first draft of the manuscript was written by Tetsu Tomonari, Yasushi Sato, and Tetsuji Takayama and all authors commented on previous versions of the manuscript. All authors read and approved the final manuscript.

## Acknowledgements

Not applicable

## Supplementary information captions

**Additional file 1: Figure S1.** Progression free survival among patients with unresectable hepatocellular carcinoma treated with lenvatinib.

**Additional file 2: Figure S2.** Overall survival among patients with unresectable hepatocellular carcinoma treated with lenvatinib.

**Additional file 3: Figure S3.** Schematic representation of the comprehensive protein phosphorylation array method.

**Additional file 4: Figure S4.** Heatmap of array proteins per cluster with significantly altered degree of phosphorylation in PLC/PRF5 cells or R2 cells treated with lenvatinib.

**Additional file 5: Table S1.** Clinical response of 53 patients with unresectable hepatocellular carcinoma treated with lenvatinb.

